# Mobile Learning for English Language Learning Assessment and Evaluation: A Review

**DOI:** 10.1101/512186

**Authors:** Monther M. Elaish, Liyana Shuib

## Abstract

Currently, one of the dominating languages in this world is English language as it has an enormous impact in practically every area of work. In this world, English language is being regarded as the language which is most commonly spoken and it is an international language for business purposes and the language of the internet which covers about 50% of the internet content. At the same time, mobile technology is becoming more and more prominent, and practitioners as well as researchers in education perceive mobile learning as a wonderful educational tool which may promote learning among students who are studying English as a foreign language. As there is a gap in literature concerning the lack of sufficient research studies which collimate their focus on the latest mobile learning technologies instead of English learning, this review intends to fill this gap. Moreover, this review orients its focus on the research problem, the acquisition of English language skills and the level of study of the participants and also it will provide a concise summary of the previous research studies by taking into consideration of group size, the type of assessment which was adopted and the duration of the experimental study in question. Therefore, this review is going to provide to the readers a thorough analysis of all the existing literature from the year 2010 to 2017 pertaining the utilisation of mobile technologies in order to study English language. This review paper focuses mainly on some key aspects that are the number of participants to be employed in such English language study, the duration of the study as well as the type of assessment and also the integration of all these factors. The result of this review can assist researchers in education sector to design accrue and fit experimental design which mitigate the challenges and facilitate the effective use of mobile learning in English language learning.

## Introduction

According to Shadiev, Hwang [1], the most widely used language is none other than English language. Therefore, it is deemed important to be proficient in English to maintain at the top level internationally. This is why so many countries put emphasis on the importance of English as a foreign language (EFL) and also to promote the awareness of English to sensitize EFL learning at various levels. In addition to that, travel and commerce as well as academic activities put a lot of attention on English which has obviously become the most importance foreign language among many non-English speaking countries [2, 3].

Portable devices including small mobile devices that can perform computing are accessible to the general public as from 1990s and these devices have gained more popularity and have experienced in computing performance owing to faster communication and technology with greater coverage in the 2000s. the industry pursues its technological evolution in order to meet the criteria of the general public for internet access learning and productivity as well as communication[4]. According to Hsu [5], for the past ten years, mobile learning (m-learning) has become the centre of attraction for practitioners, as well as researchers, because of the fast evolution of the sector of telecommunication technology and its related application to portable devices. These devices comprise of smart phones, mobile phones, Tablet PCs and personal digital assistants (PDAs). M-learning is recognised as greatly helpful tool in many sectors such as educational sector and educational scenarios that include independent learning, collaborative learning, and lifelong learning. English as a foreign language (EFL) learning is no exception [6].

The utilisation of these mobile technologies appeared to be in line with the strategic educational aims that are the improvement of the student retention and achievement, and the support of the difference of the various learning needs as well as to reach learners who would not have the chance to be participants in education.[6]. A lot of energy has been spent to understand how the mobile technologies are related to the traditional and the innovative methods of learning and teaching and to demonstrated the applicability of the m-learning across the wide range of activities [7, 8] and to pinpoint the most crucial emerging problems [9, 10].

According to Alghamdi and Shah [11], a review of mobile-assisted language learning (MALL) research as from the year 1994 to 2012 revealed more than 500 MALL research scientific studies. The mobile-assisted language learning (MALL) is a method learning a language whereby a handheld mobile device is used to assist and enhance learning. As smart mobile phones and other types of hand-held devices are ever increasingly developed, it is thought that their integration in the educational sector as well as in EFL classrooms will always rise. According to Crompton and Burke [12], research in mobile learning has been divided and is closely related to the understanding of the experimenter. Moreover, there is still few systematic information which is available based on over 20 years of mobile learning research. Next, a specific type of study which is the descriptive/diagnostic study is being looked at. In both diagnostic and descriptive studies, the researcher should define clearly vividly what measurements he wants to take he wants to measure and search for suitable techniques.

In descriptive/diagnostic research study the first action is to mention the aims and objectives with much precision in order to make sure that the collected data are all relevant. Since there are various types of methods for data collection, the researcher must be able to select well the method of his study by keeping in mind the following factors that are : (1) Availability of funds, (2). Nature, scope and object of enquiry (3). Precision required, (4) Time factor. Each method has its own specific uses and one must be aware of when to use and in which situation. The most desirable methodology to be employed depends on the amount of money available to conduct this research and the nature of the problem as well as the time and resources and the level of accuracy which is needed. However, having mentioned all this, the ability and experience of the researcher is very important. [13].

Based on the viewpoints of Webster and Watson [14], a literature review is an important characteristics of any academic project as it creates a firm base to advance technical knowledge. It makes the development of theory smooth, shuts down areas where an abundance of research exists, and searches areas where research is required. In this research, a quasi-experimental research design was employed to investigate the effectiveness of the mobile technology within a collaborative learning experience to ease the engagement of the students and the critical thinking. Through the use of a multi model technique of analysing effectiveness according to some research study, myriads of types of data sources were utilised. These types of data include student questionnaires, completed written product, as well as classroom behavioural observation [15].

According to Wingkvist and Ericsson [16], research techniques are crucial parts of how the research is being conducted.. Both research evaluations as well as basic research are imperative as they support research field to grow up and also researchers to avoid making the same mistakes. As mobile learning matures, it is essential to investigate how this line of research is being performed. In the meanwhile, one needs to understand the impact of technology and also the knowledge which is produced. As a consequence, this creates challenges to all facets of the mobile learning research. The techniques that are considered are mainly case studies, field studies, experimental studies, survey research, basis research and action research and well designed and executed experimental studies. All of them can be replicable and this eases data collection. In mobile learning research, experimental research studies are appropriate for the evaluation of design ideas, specific products and theories about the design and the interaction between users in controlled environmental conditions without interference from the outside world. Field studies and experimental studies are the most common types of research studies. By definition, 100% of the artificial research used experiment research.

Although there are a lot of qualitative analyses of the application of mobile devices in educational sector, there is a still lack of systematic quantitative analyses of the effects of mobile-integrated education. This experimental study conducted a metaanalysis and research synthesis of the effects of integrated mobile devices in teaching and learning. It was found that 110 experimental and quasi-experimental journal articles that were published as from the year 1993 to 2013. These research studies were coded and analysed. In general it was found that there was a moderate mean effect size of 0.523 for the usage of the mobile application to educational sector. The effect size of the moderator variables were assessed and the disadvantages as well as the advantages of mobile learning in different levels of the moderator variables were also assessed based on the content analysis of the individual studies [17].

Ok and Ratliffe [4] also discovered that there were 4 review articles that were published between the year 2010 and the year 2015 that investigates research based on m-learning and MALL. These reviews consist of 289 research studies and also identifies common weaknesses that a lack of theoretical frameworks, the quality of the research studies were also poor and they were excluded from the reviews and also it was noticed that there was a lack of quantitative studies on outcomes and subjective deductions.

Duman, Orhon [18], analysed how the MALL has been evolving throughout the recent years and its features. These research studies published in international reviews were listed in the Social Sciences Citation Index (SSCI). The sixty-nine studies that fit the suggested time frame and the study parameters were investigated using a classification form. The results demonstrate that research in the field has rocketed from 2008 and reached its peak in 2012. Since then, the teaching of vocabulary with the use of cell phones and PDAs has remained popular. A lot of studies did not relate their research on any theoretical framework. As a matter of fact, applied and design-based research was prominent in the field, and these studies adopted quantitative research techniques. After analysing these results, future directions were suggested as well as future practices that need to be adopted in the field. All the review studies which have been examined in this research did not specify any concrete assessment and evaluation method which is suitable, also they did not assess the appropriate group size of participants for this particular type of project and the subgroups or groups needed and there is also a missing assessment about the duration of the research study especially related to experimental study using mobile learning for English language learning. However, Elaish, Shuib [19], Elaish, Shuib [20] have reviewed currently the utilisation of mobile English language learning and suggested some common assessment and evaluation ways in this area of research. They have also reviewed the various points that are as the suitable type of assessment to be used for an experimental study, the size of groups and the duration of the experiment.

Generally, many research studies investigate the facets of m-learning for both the researchers as well as the instructional designers and also pays attention on the effective usage of the latest mobile learning technologies for education [21], but did not focus on English language learning. On the contrary, there are many review studies on mobiles and ubiquitous learning [12, 21-28]. Therefore, this review focuses on three factors ((i) research problem, (ii) participants’ level of study, (iii) English skills) and their effect on groups’ size, the duration of the experiment and which assessment(s) is appropriate and also fits for the study. It is worth to note that all the mentioned studies about the duration of the research study, the groups’ size and the type of the assessment to be used. This research study endeavours to provide a comprehensive analysis of the current studies to understand the assessment and evaluation of mobile English language learning. The findings which emanate from this study are expected to provide an expansion to the existing body of knowledge on the learning of English language to help users, researchers as well as policy makes and practitioners in the educational sector in their allocation of required resources and also to make plans to support upcoming researches and application, and hence improve educational practise. While performing any experiment, it is important to know the suitable size of the number of participants to be recruited so as on can draw statistical inferences from these findings. Also the duration of the experimental research is crucial so as the students are being provided with the required amount of time for them to absorb this type of new mobile learning technology. Each assessed variable will help everyone including future researchers to understand the effect of the mobile application technology in terms of motivation as well as one need to be aware of the right statistical method to assess each measured variable.

### Research Questions

- What is the groups’ size of the experiment for each factor?
- What is the duration of study for each factor?
- What is the suitable and fit assessment for each factor?
- Does the result from Q1-Q3 questions is used on real studies when combine all the factors and which one has the power to effect on the researchers decision?

## Method

Most systematic reviews obey 5 key rules that are (1) framing the question, (2) identifying relevant publications, (3) assessing the study quality, (4) summarising the evidence and (5) interpreting the findings [29]. A review possesses the adjective systematic if it is based on a clearly calculated question, and also it involves the relevant identification of studies, quality appraisal and also the summary of evidences through the use of explicit methodology.

The review is based on author name [19] study which encompasses the recent systematic review studies of utilising mobile learning for English language learning. A review of the articles which published between the year 2010 and 2017 was conducted. Our search was restricted to research articles on the role of m-learning applications in enhancing English language learning using a systematic search technique as shown in Figure 1.

**Figure 1:**
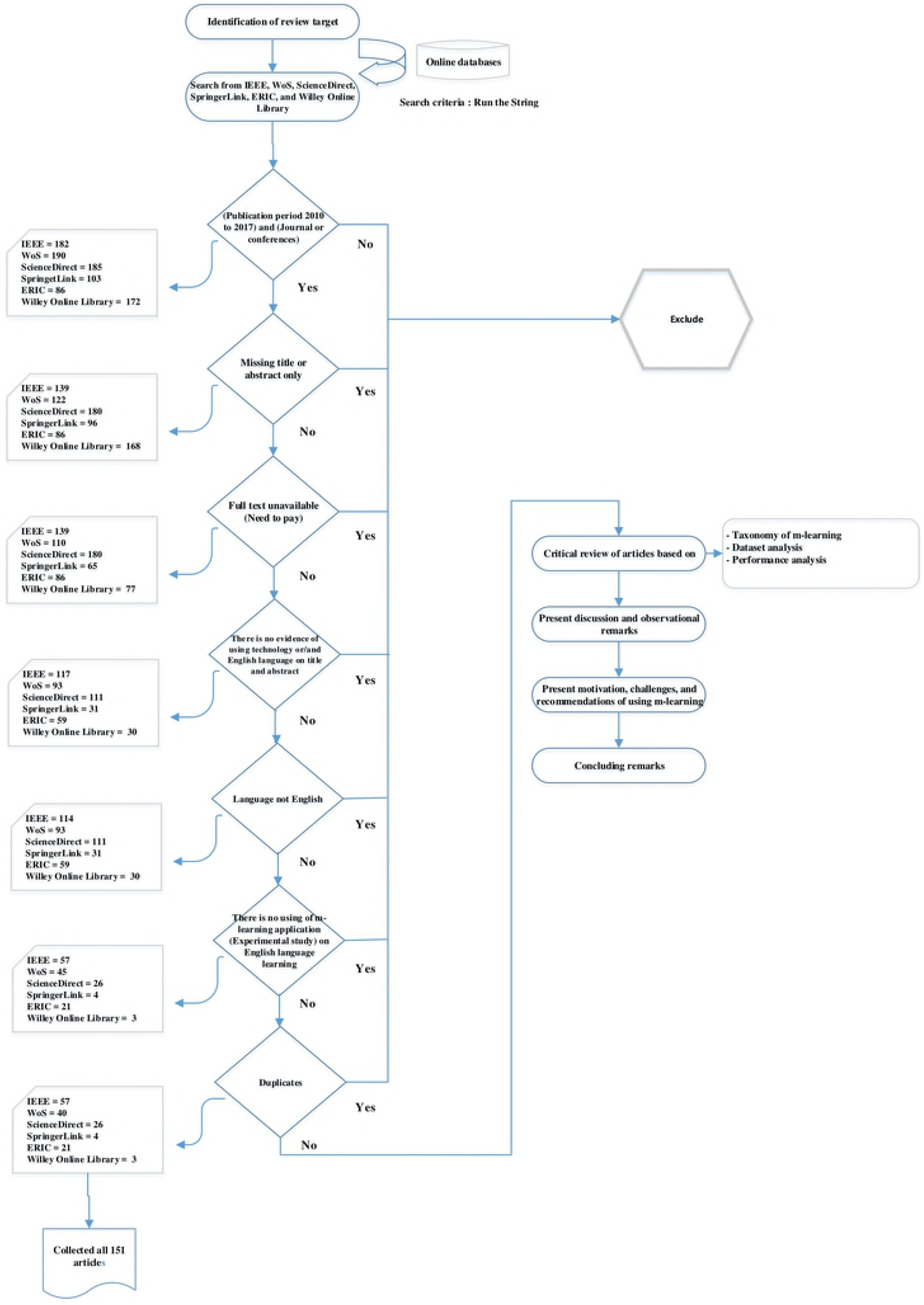
Flowchart of the study steps.

### A. Review protocol

A review research protocol was created to analyse the proposed research questions, search strategy, the selection of databases, exclusion and inclusion criteria, and assessment criteria. The process followed the steps described in Figure 1.

#### 1) Research question

In order to perform a systematic review study, the main research question had to be created first and after the research questions have been specified, the review procedure involved building search strategies to recognize and extract relevant studies. The main intention of the review was to identify the duration of the study, group size, the assessment and evaluation in using m-learning for English language learning for English language problem, the participant, and English language skills, and to provide a suitable way to select the fit method when these factors are together in one study.

#### 2) Search Strategy

In order to develop an effective search strategy, keywords were identified together with alternative methods of expressing them. Fellow academics were consulted to make sure that all the possible alternatives were covered. The search string to ‘_t’ was altered as it represents the formatting requirements for each database. The general strategy was as follows: “mobile learning OR m-learning OR mlearning OR personalized learning OR ubiquitous learning OR u-learning OR anytime and anywhere learning OR mobil* learn* AND English language” [19].

#### 3) Literature databases

The primary search was conducted from following databases (Web of Science (Wos), ScienceDirect, IEEE Xplore, Wiley Online Library, SpringerLink, Education Resources Information Center (ERIC)). All the available educational databases were searched to select a technical literature only.

#### 4) Study exclusion/inclusion criteria

To ensure that the most relevant researches were included, the publication period was restricted from January 2010 to December 2017. Only those articles that were written in English language were included. The criteria below were used to determine if the article would be included in this review:

- The articles were published between 2010 and 2017
- The full text was available (i.e. conference abstracts were excluded);
- The topic was English language acquisition through m-learning;
- The articles were written in English language.

### B. Selection Process

The following process represents the steps that were applied in order to conduct the research:

#### 1) Conferences and journals studies

During the first step, all unpublished conference and journal studies were excluded manually.

#### 2) Full-text availability

In the next step, any research articles for which the completed text was not available was excluded. This included the conference abstracts, letters addressed to editors and advertisements.

#### 3) Full-text unavailable

In the third step, any article which is not accessible from the available data sources was not included.

#### 4) Use of technology for English language acquisition

In the fourth step, if there was any evidence of using/investing of technology for English language learning from the articles’ titles or abstracts then they was excluded.

#### 5) Non-English written studies

In the fifth step, any articles that were no written in English language were excluded.

#### 6) Use of m-learning application for English language learning

In the sixth step, after finish reading of the included articles (full-texts), those that did not show any evidence of using m-learning application for English language learning were excluded.

#### 7) Duplicates

In the last step, any duplicated articles were excluded.

### C. Data report process

Lastly, all research articles were summarized and their data were entered into tables, including author(s) names and year of publication, English language problems, participants, assessment, English language skills, duration of the study, group size. These items were selected based on the objective and research questions of this study. After this step has been done, the analysis of these data were started to answer each of the research questions.

## Findings

### The groups’ size of the experimental for each factor

This component of this research investigates which group size is better for (English problem, participant, and English skills). Tables in this section are shown the groups size for each English problem, types of participant, and English skills. Moreover, the group size could be divided to three categories (one group (non-groups), groups, non-mentioned). 54 of 151 of the analysed articles are non-grouped in which there was only one group for the experimental, 68 of 151 have divided participants into groups (two groups or categories in most of the cases), and 19 of 151 have not mentioned anything about participant and how their studies dealt with them to do the experimental.

### English problems

English language problems are divided into the group size categories. Every English language problem has different group size as: the best group size for Lack of identify needing, reports, or studies or testing the effect of technologies (14 of 151 articles) is from 100 and above without divided participants into groups, and motivation problem studies can divide participants into groups (22 of 151) from 30 to less than 99. These two English problems are the most English problem appeared on the studies (95 of 151). The rest is shown in the Table 1. For ease of denotations (Labelling in Table 1), let A represents 30 > x, B represents 30 =< x < 99, C represents 100 =< x <= 6000, D represents 100 =< x =< 240 and E represents none.

**Table 1:** Group size based on English problem.

| English Problems | Group Size |  |  |  |  |  |  |
| --- | --- | --- | --- | --- | --- | --- | --- |
|  | Non-Groups |  |  | Groups |  |  |  |
|  | A | B | C | A | B | D | E |
| Lack of the identification of reports, or studies or testing the effect of technologies (L) | 4 | 6 | 14 | 1 | 9 | 4 | 3 |
| Motivation (M) | 9 | 6 | 9 | 1 | 22 | 3 | 4 |
| Other (O) | 6 | 1 | 3 | 2 | 4 | 3 | 3 |
| Language difficulties or Limitation of vocabulary and the reading materials is not available (LD) | 2 | 0 | 1 | 0 | 6 | 6 | 4 |
| Lack of equipment or poor current equipment, technique, or software (LQ) | 0 | 0 | 1 | 0 | 1 | 0 | 5 |
| Culture (C) | 0 | 0 | 1 | 1 | 1 | 0 | 0 |
| Unstandardized curriculum (U) | 0 | 0 | 1 | 0 | 3 | 0 | 0 |
| Unqualified English teachers or improving their quality (UQ) | 0 | 0 | 0 | 0 | 1 | 0 | 0 |
| TOTAL | 21 | 13 | 30 | 5 | 47 | 16 | 19 |

### Participants

As shown in Table 2, 19 of 151 studies did not show any grouped or mentioned about participants. However, university and school students and learners were employed without mentioning their levels (115 of 151). Moreover, some of them show that grouped students from 30 to less 99 are the best way to do the experiment for this kind of study. The group size based on the participants are summarised in the following Table 2. For ease of denotations (Labelling in Table 2), let A represents 30 > x, B represents 30 =< x < 99, C represents 100 =< x <= 6000, D represents 100 =< x =< 240 and E represents none.

**Table 2:** Group size based on participants.

|  | Group Size |  |
| --- | --- | --- |
|  | Non-Groups | Groups |

| Participants | A | B | C | A | B | D | E |
| --- | --- | --- | --- | --- | --- | --- | --- |
| University students (Univ) | 10 | 6 | 13 | 0 | 19 | 10 | 19 |
| School/pre-school students (S) | 0 | 0 | 1 | 0 | 15 | 3 |  |
| Learners/users/students(not specific) (NS) | 8 | 5 | 9 | 3 | 11 | 2 |  |
| Mixed: students and teachers (school/university) (S+T) | 3 | 0 | 6 | 2 | 1 | 0 |  |
| Teachers (T) | 0 | 2 | 0 | 0 | 0 | 0 |  |
| Mixed: school and university students (S+US) | 0 | 0 | 0 | 0 | 0 | 1 |  |
| Others (O) | 0 | 0 | 1 | 0 | 1 | 0 |  |
| **TOTAL** | <u>21</u> | <u>13</u> | <u>30</u> | <u>5</u> | <u>47</u> | <u>16</u> | <u>19</u> |

### English skills

The studies which studied all the English skills (or do not mentioned about which skills they studied) put all the participants in one group which its size from 100 and above. Speaking skill is in the same with all English skills. On other hand, divided participants into groups from 30 to less 99 is the best way to study vocabulary which ranks on the second position behand all English skills. In additional, writing, reading, and more than 1 skill and less than all are in the same line with vocabulary. For ease of denotations (Labelling in Table 3), let A represents 30 > x, B represents 30 =< x < 99, C represents 100 =< x <= 97746, D represents 100 =< x =< 240 and E represents none.

**Table 3:** Group size based on English skills.

| English Skills | Group Size |  |  |  |  |  |  |
| --- | --- | --- | --- | --- | --- | --- | --- |
|  | Non-Groups |  |  | Groups |  |  |  |
|  | A | B | C | A | B | D | E |
| All | 9 | 7 | 17 | 4 | 10 | 3 | 19 |
| Vocab (V) | 2 | 2 | 2 | 0 | 15 | 7 |  |
| Writing (W) | 0 | 1 | 0 | 0 | 3 | 0 |  |
| Reading (R) | 1 | 1 | 2 | 0 | 8 | 3 |  |
| Listening (Li) | 2 | 0 | 0 | 0 | 2 | 3 |  |
| Speaking (S) | 4 | 0 | 6 | 0 | 3 | 0 |  |
| Grammar (G) | 0 | 0 | 2 | 0 | 0 | 0 |  |
| More than 1 and less than all (2 and M) | 3 | 2 | 1 | 1 | 6 | 0 |  |
| **TOTAL** | <u>21</u> | <u>13</u> | <u>30</u> | <u>5</u> | <u>47</u> | <u>16</u> | <u>19</u> |

## Duration of study for each factor

This part is going to find which the duration is better for (English problem, participants, and English skills). Tables in this section are shown the duration for each English problem, participants, and English skills. Moreover, the duration could be divided into many categories. 52 of 151 have not mentioned any duration for the experiment and the others classified in many categories.

### English Problems

The study duration from 1 month to 4 months (1 semester) is the best duration to study the lack of identification of, reports, or studies or testing the effect of technologies. However, lack of motivation can be studied in three categories (1 week to 4 weeks, 1 month to less than 4 months, and 4 months to less than 12 months (1 academic year)) but 1 week to 4 weeks is the best as shown in Table 4. For ease of denotations (Labelling in Table 4), let A represents 1 week>x, B represents weeks, C represents months, D represents months, E represents None, F represents 1 year =<x, and G denotes other 16 non-formal sessions, 3 sessions among study time.

**Table 4:** Duration based on English problems.

| English Problems | Duration |  |  |  |  |  |  |
| --- | --- | --- | --- | --- | --- | --- | --- |
|  | A | B | C | D | E | F | G |
| Lack of identify needing, reports, or studies or testing the effect of technologies | 0 | 3 | 10 | 5 | 52 | 7 | 4 |
| Motivation | 1 | 15 | 12 | 11 |  | 1 |  |
| Other | 2 | 3 | 6 | 1 |  | 3 |  |
| Language difficulties or Limitation of vocab and reading materials is not available | 1 | 4 | 3 | 3 |  | 1 |  |
| Lack of equipment or poor current equipment, technique, or software | 0 | 0 | 0 | 1 |  | 0 |  |
| culture | 0 | 0 | 0 | 0 |  | 0 |  |
| unstandaridized curriculum | 0 | 0 | 1 | 0 |  | 0 |  |
| Unqualified English teachers or improving their quality | 0 | 0 | 1 | 0 |  | 0 |  |
| **TOTAL** | <u>4</u> | <u>25</u> | <u>33</u> | <u>21</u> | <u>52</u> | <u>12</u> | <u>4</u> |

### Participants

Table 5 shows that studied which focus on university students is held from 1 month to less 4 months or 4 months to less 12 months (is the best). School students are trained in 1 week to 4 weeks or 1 month to less 4 months. In this line with school students, learners also show same but 1 month to less 4 months is the best. The following denotations are used to describe the various columns of the following table and they are as follows. A denotes 1 week>x, B denotes 1 week =< x< =4 weeks, C denotes 1 month < x< 4 months, D denotes 4 months <= x < 12 months, E denotes None, F denotes 1 year =<x and G denotes Other 16 non-formal sessions, 3 sessions among study time.

**Table 5:** Duration based on participants.

| Participants | Duration |  |  |  |  |  |  |
| --- | --- | --- | --- | --- | --- | --- | --- |
|  | A | B | C | D | E | F | G |
| University students | 2 | 8 | 12 | 13 | 52 | 6 | 4 |
| School/pre-school students | 0 | 7 | 7 | 0 |  | 2 |  |
| Learners/users/students(not specific) | 2 | 8 | 11 | 4 |  | 2 |  |
| Mixed: students and teachers (school/university) | 0 | 2 | 3 | 2 |  | 1 |  |
| Teachers | 0 | 0 | 0 | 1 |  | 1 |  |
| Mixed: school and university students | 0 | 0 | 0 | 1 |  | 0 |  |
| Others | 0 | 0 | 0 | 0 |  | 0 |  |
|  | <u>4</u> | <u>25</u> | <u>33</u> | <u>21</u> | <u>52</u> | <u>12</u> | <u>4</u> |

### English skills

Table 6 shows that all English skills studies (13 of 151) have done between 4 months to less 12 months. However, vocabulary studies (10 of 151) have done between 1 week to 4 weeks and reading (5 of 151) studies also have done in the same duration. All these English skills are shown as the most skills appeared on the studies (71 of 151). The following denotations is used to describe the various columns of the following Table 6 and they are as follows. A denotes 1 week>x, B denotes 1 week =< x< =4 weeks, C denotes 1 month < x< 4 months, D denotes 4 months <= x < 12 months, E denotes None, F denotes 1 year =<x and G denotes other 16 non-formal sessions, 3 sessions among study time.

**Table 6:**
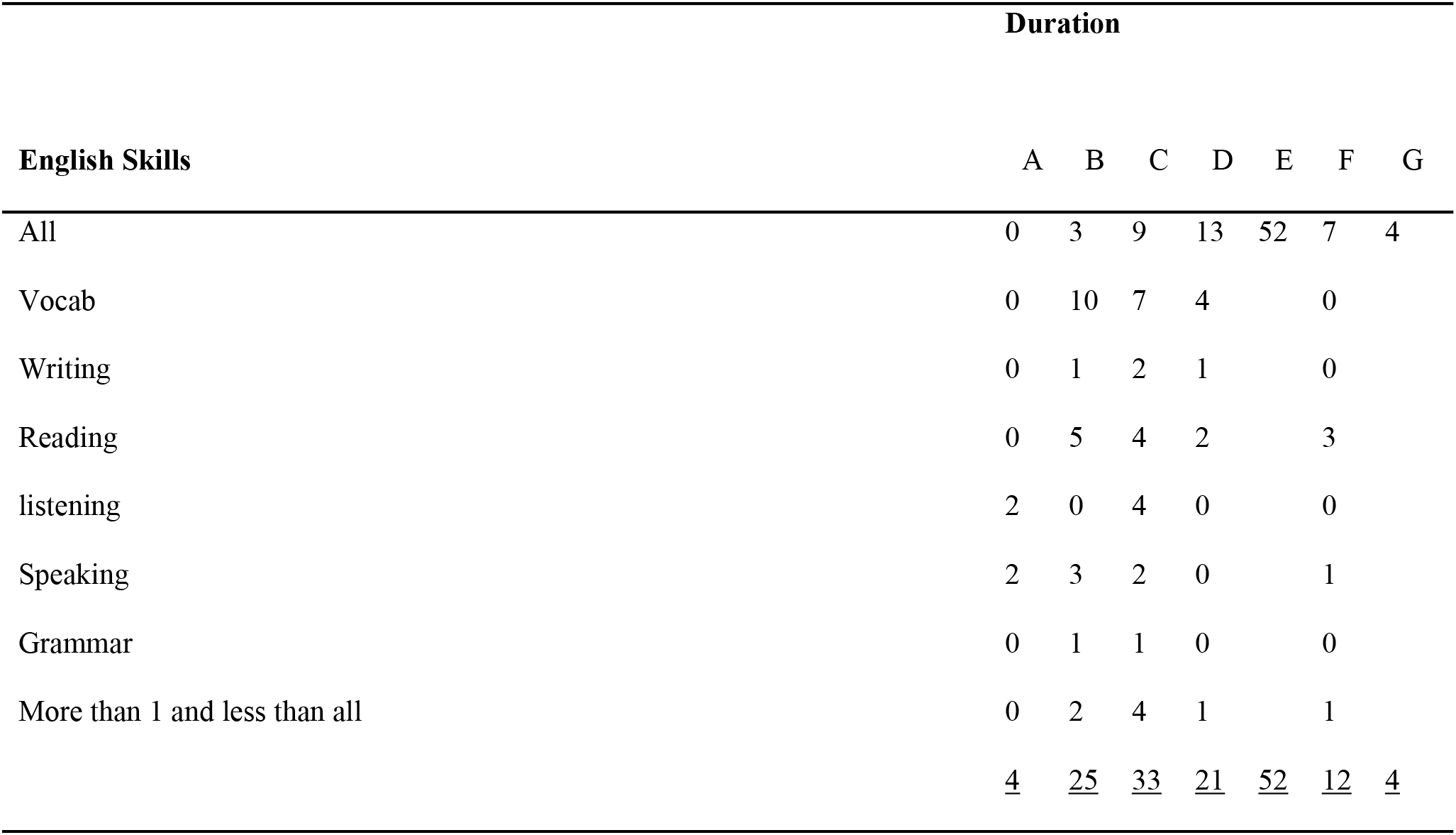
Duration based on English skills.

## Suitable and fit assessment for each factor

This part is going to find which the assessment is better for (English problem, participants, and English skills). Tables in this section are shown the assessment for each English problem, participants, and English skills. Moreover, the assessment could be divided into many categories. 9 of 151 have not mentioned any assessment for the experiment and the other classified in many categories.

### English Problems

Questionnaire is the most used assessment for lack of identify needing, reports, or studies or testing the effect of technologies studies. In other hand, test and questionnaire is the assessment mostly used for lack of motivation and Language difficulties or Limitation of vocabulary and reading materials is not available as shown in Table 7.

**Table 7:**
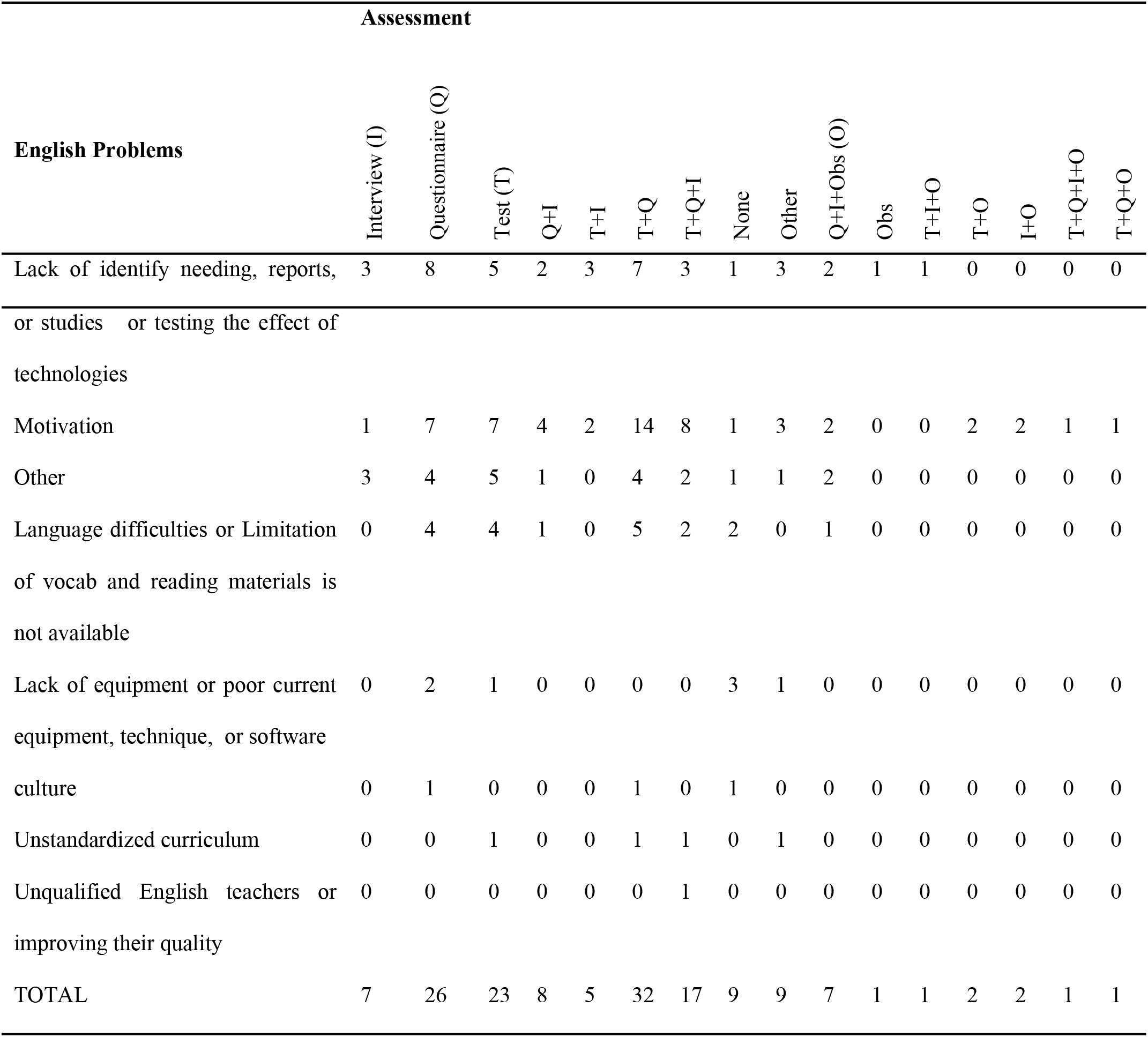
English problem based on assessment.

### Participants

Table 8 shows that the studies which focus on university (18 of 151) and school (7 of 151) students have used test and questionnaire as assessment method plus test, questionnaire, and interview as same as assessment method for students school. But learners’ studies have used test as assessment method.

**Table 8:**
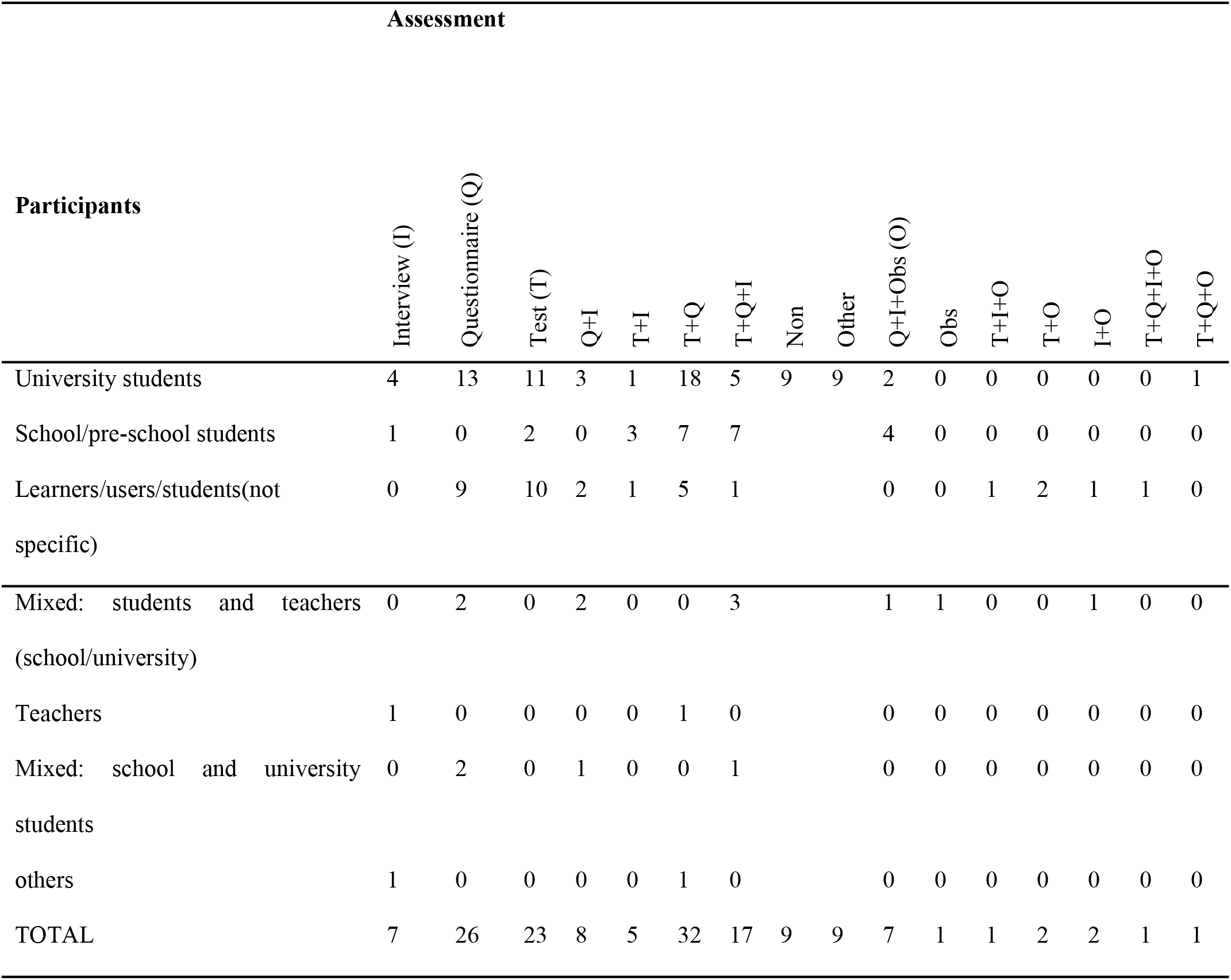
Participants based on assessment.

### English Skills

Questionnaire (12 of 151) is the mostly used assessment method for all English skills. Only 5 research studies use speaking as a method of assessment (5 of 151) based on questionnaire. On other hand, test assessment comprises of (25 of 151) which 9 of them focus on vocabulary as shown on Table 9.

**Table 9:**
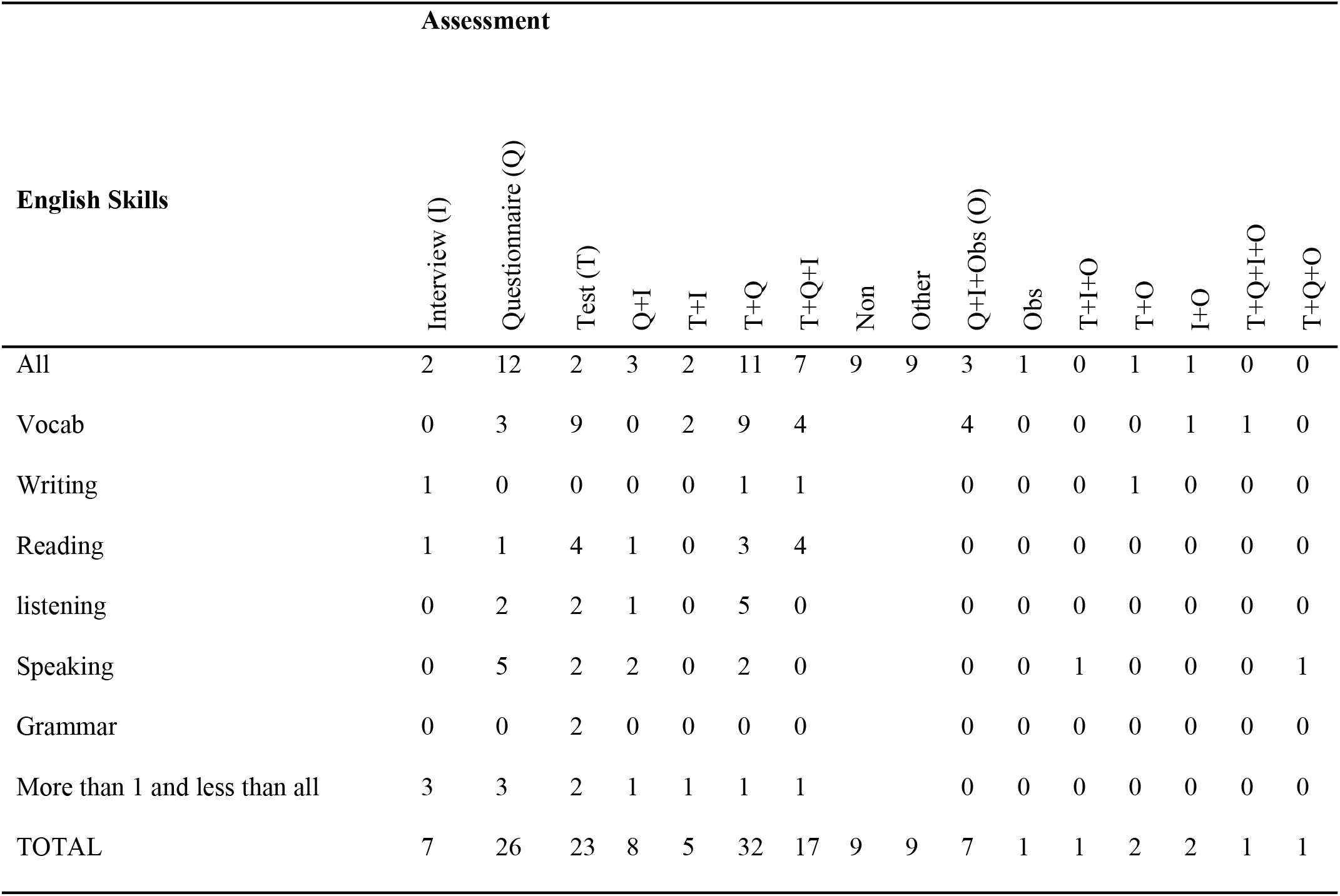
English skills based on assessment.

## Pattern to select the groups’ size, duration of the study, assessment method for the study when take into account the three factors (English problems, participants, English skills) together

To answer this question Table 10 has prepared based on three things: First, Web of Science database articles because it is interdisciplinary and covers all scientific areas[30, 31], but it only covers what it considers to “best” journals and concentrates on English language ones [32]. Additionally, it is a curated collection of over 20,000 peer-reviewed; high-quality scholarly journals published worldwide (including Open Access journals) in over 250 science, social sciences, and humanities disciplines. Conference proceedings and book data are also available [33]. Additionally, WoS remains an indispensable citation database [34]. second, the studies which showed no missing for any factor so any study does not show clearly the groups’ size, duration of the study, or assessment method are omitted. Finally, cannot use all the studies because their large number which cannot be shown on the single article.

**Table 10:**
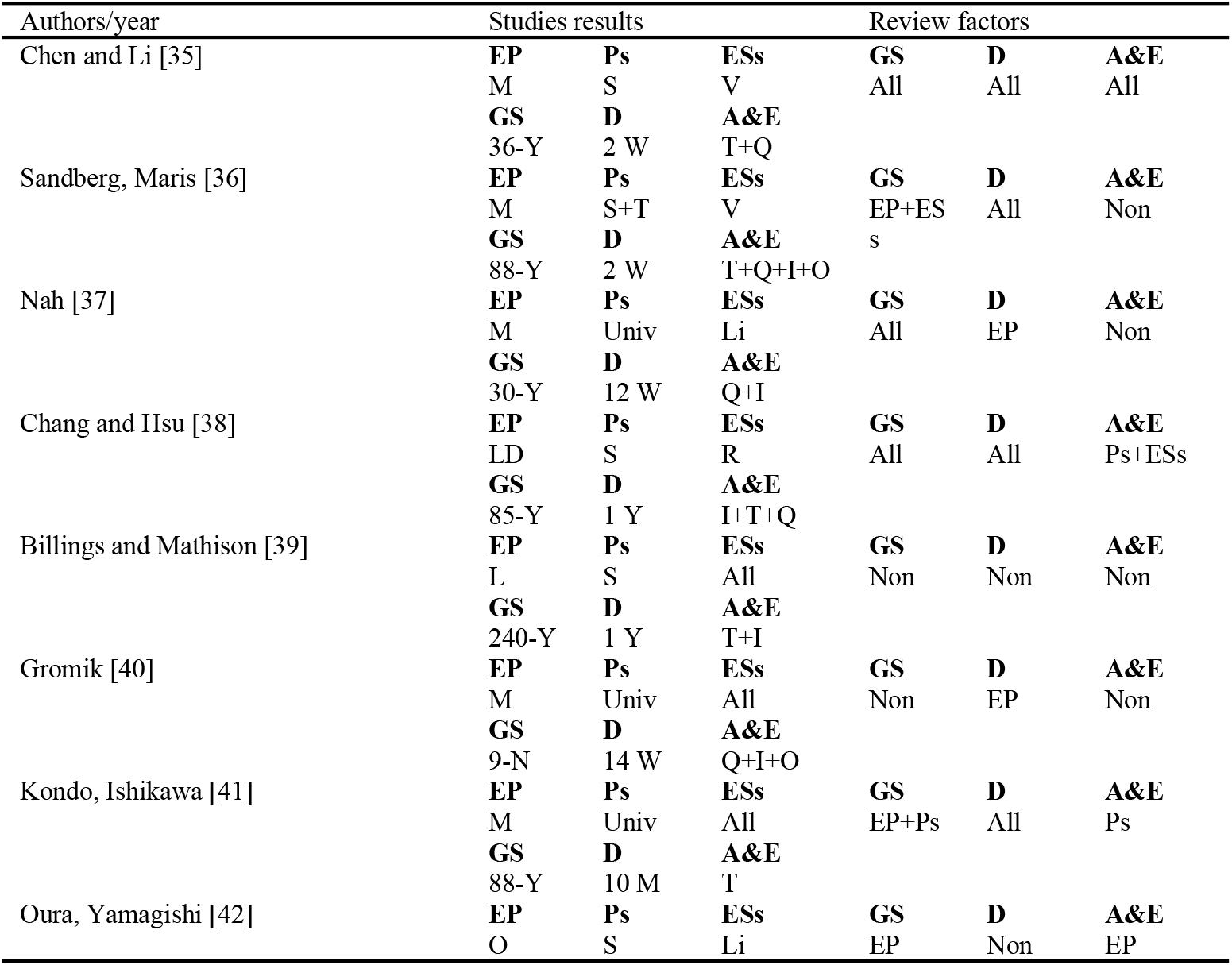
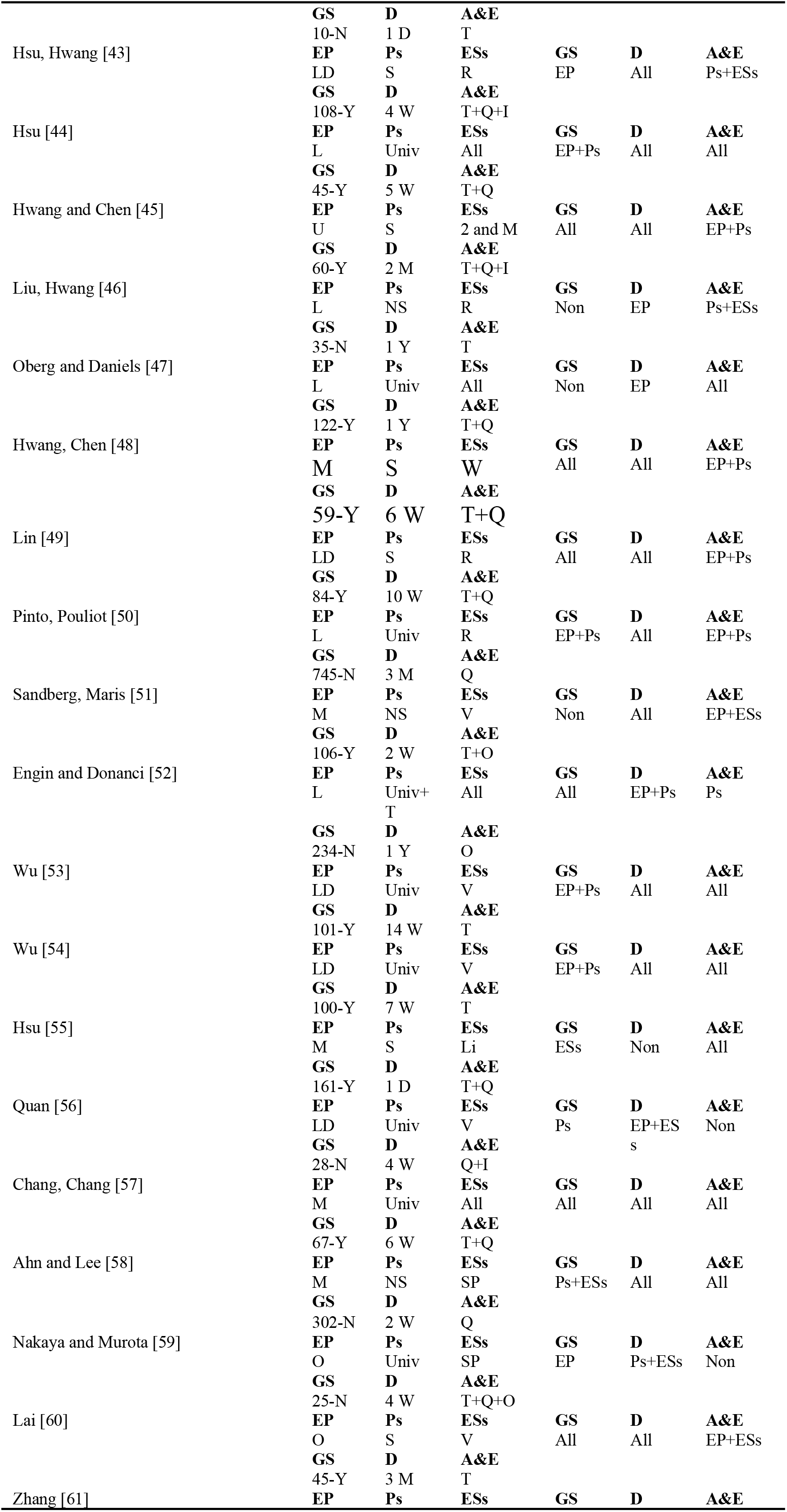
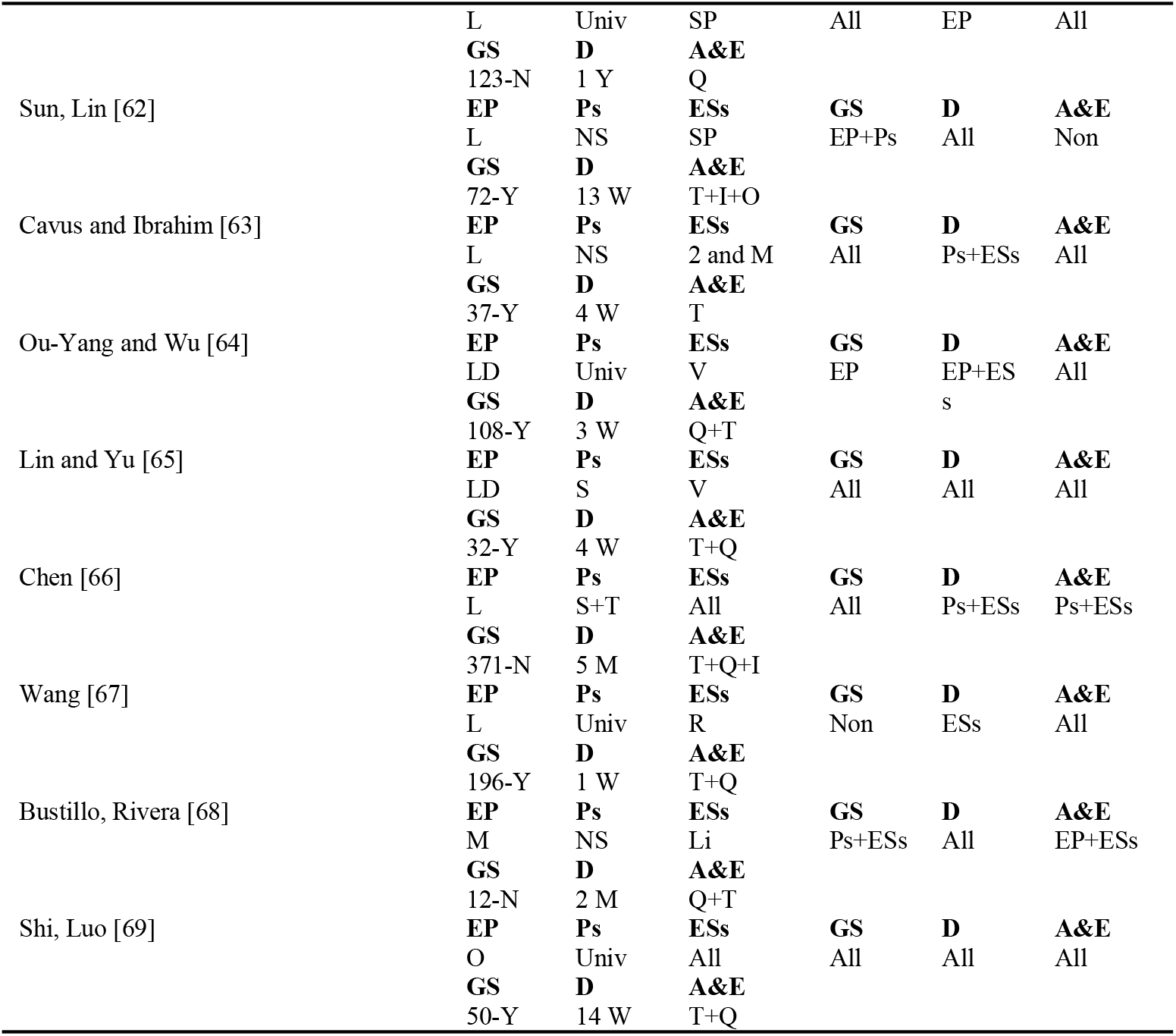
Web of Science studies (the real results and the supposed results)

The real studies finding has been addressed on the table plus the suitable finding from the three research questions as well. Then the study has pointed whether each study follow the review findings or not and is there any pattern (which factor gets the power to be selected by the researchers) to selected the factors. On the tables below studies results column shows what the researchers have chosen to (groups’ size (GS), duration (D), assessment and evaluation (A&E)). The review factor column shows which factors (English problem (EP), participants (Ps), English skills (ESs)) have the power that effect on the researchers’ decision to select the GS, D, and A&E.

## Discussion

This section discusses the results in terms of the groups’ size, participants and assessment and evaluation. As first instance, the research question RQ1 is addressed which related to the experimental for each factor. 45% of the studies out of the total of 151 have divided the students into more than 2 groups. However, almost 35.7% of these studies did not use grouping as a factor to divide students into subgroups. According to Price, Jhangiani [70], in the context of one-group design, a processing is implemented (or an independent variable is manipulated), and then a dependent variable is measured once after the implementation of the treatment. For example, a researcher is interested in the effectiveness of an anti-drug education program on the attitudes of elementary students’ school to illicit drugs. The researcher could implement the drug control program, and immediately after the end of the program, measure students’ attitudes toward illicit drugs. This is the weakest type of quasi-experimental design. The absence of a control or comparison group is a major drawback of this design. There is no way to determine what would have been the attitude of these students if they had not followed the drug program. About 1/3 of the studies did not mention about the accuracy of their results as the experimental design was incorrect.

For grouped studies, the study has investigated which the suitable number of participants for the groups for each factor. The range 30 =< x < 99 is the good number of participants which should be divided into groups. 47 of 151 (47 of 68 of grouped studies) studies have used this number of participants in their studies. In details, for English language factor: Motivation showed the highest number of studies (22 of 47) which have selected this range of participants for their studies then follow by Lank of the Identification (9 of 47). Regarding to participants type university students showed the highest number (19 of 47) however, school students ranked closely to it (15 of 47). Finally, vocabulary skills came firstly by 15 of 47 after that all English skills 10 of 47. In non-grouped studies, 30 of 151 of studies (30 of 65) have selected 100=< x to participant them in one group. But 30> x has also scored a second highest number (21 of 65) which is not far from the top. So in this case it is too hard to find a suitable pattern in general plus this way is not a good way for run experimental study. So the study is going to focus on each factor independently.

The second factor that will be discussed here concerns the duration of study. About 34% of the research studies have not shown any duration of their studies which addressed as a main gap of these studies due to the important of the duration of study to determent whether the study gave a enough time to the participants to train on the app and understand its propose and functions. Key [71], have listed out steps to conduct the experimental study which one of them determine place, time, and duration of the experiment. However, there is no specific duration of this kind of study which can be used. 99 of 151 have showed their period of the study which there is there periods can be used on this kind of studies. 1 months < x < 4 months is the highest one (33 of 99) then 1 week =< x <= 4 weeks (25 of 99) then 4 months <= x < 12 months (21 of 99).

In details, 1 months < x < 4 months is the best study period for Lack of Identify English problem. On other hand, motivation studies have been occurred 1 week =< x <= 4 weeks (15 of 99) as the best study period. In additional, 1 months < x < 4 months (12 of 99) and 4 months <= x < 12 months (11 of 99) are shown as second and third duration of motivation studies. For participants type, students university have been participated on studies which have contented 1 months < x < 4 months (12 of 99) and 4 months <= x < 12 months (13 of 99) however, for school students 7 of 99 have done 1 week =< x <= 4 weeks and 1 months < x < 4 months. Additionally, learners without specific have been participated on studies which have occurred 1 week =< x <= 4 weeks (8 of 99) and 1 months < x < 4 months (11 of 99). Regarding to English skills, 4 months <= x < 12 months (13 of 99) is the best period for all English skills studies. 1 week =< x <= 4 weeks is the best for vocabulary skills one. In this section, the period of study is very because there are different period based on the factors and their categories.

The third research question (RQ3) concerns about the suitable and the fit assessment. In this kind of research most of the researchers have been mentioned that they used experimental research. According to Klazema [72], Sukamolson [73], experimental research is type of quantitative research. Even though, there are 81 of 151studies have used quantitative research only but there are 44 of 151 studies are mixed (quantitative and qualitative), this finding is in the line with [74] which they demonstrate the value of combining qualitative and quantitative methods. The study showed that 8 of 151 studies are qualitative research. The highest assessment is questionnaire mixed with test (32 of 151) 32 of 81 of quantitative research studies.

For English problems, questionnaire (8 of 151), and questionnaire and test (7 of 151) are the assessment methods for lack of Identify. However, motivation problem studies have used questionnaire and test (14 of 151) as assessment method. Studies which involved university students on the experimental has used questionnaire and test (18 of 151), and questionnaire (13 of 151) are the assessment method but test (11 of 151) is also used widely. Regarding to English skills, the studies which used the app to improve all the English skills have used questionnaire (12 of 151) as the assessment method. But studies which used vocabulary have used test (9 of 151) and test and questionnaire (9 of 151) as the assessment method.

9 of 151 have not mentioned any assessment method and same number have used different assessment method such as recoding. Finally questionnaire and test are the best way as assessment method for each factor whatever they are used separated or combined.

The fourth research question combines all the previous three questions especially to pinpoint studies that combine all those factors together and when to select a particular factor. The review shows that only 99 of 151 have shown together groups’ size, duration, and assessment. Table 10 has listed Web of Science studies (35 of 151) 35 of 99 studies. The table was prepared based on point of view of the authors because the review results show different factors for each aspect. So the authors tried to select the suitable and logical result for the table. The table shows that English problem (25 of 35) has the most powerful effect to select the group size so the researchers were influenced by this factor when they wanted to selected the group size of the study. As same as group size, the studies’ duration has been effected by English problem (28 of 35). On the other hand, assessment and evaluation of the studies have influenced by participants type (24 of 35) mostly.

## Conclusion

This review paper has provided a detailed meta-analysis of the contemporary experimental research studies from the year 2010 to 2017 concerning the use of mobile assisted technologies to promote the learning process of English language. It was found, for the one-group factor, that some studies did not have even a control group in order to compare their findings with which make it difficult to really validate their findings or whether the findings are biased or unbiased. For those experimental studies which employ grouping, about 31% of the total investigated studies or 69% of the grouped studies use about 30 to 98 as the number of participants in their study. For those studies which employ motivation as a factor, it was found that 22 out of the 47 studies investigating motivation selected the aforesaid range of participants (that is 30 to 98).

The second factor which this review has investigated concerns the duration of the experimental study, it was demonstrated that the range of time between 1 to 4 months displayed the highest frequency of time duration usage by the researchers and also those motivation studies did employ similar duration for their experiments which represents the best study period. The next important factor which this review paper investigated concerns the type of research and it was also shown here that more than 53.6% of the studies employed the quantitative approach only while 29.1% of the studies employed a mixed quantitative and qualitative approach. In the same line of thought, it was found that the 11.9% of the studies involving university students employed test and questionnaire while those studies which utilise the mob app to improve all the English language skills represented only 7.9%. In addition, those studies which employ test to check the vocabulary was found to be 5.9% and this same figure was obtained for those studies which employ test as well as questionnaire. Last but not least, 65.5% of all the examined studies employ all these factors altogether that are the size which is the number of participants, the duration of the study as well as the type of assessment. Future reviews can include a longer period of time and also with a larger database of research articles that this research may miss as it was limited to search terms that the researchers desired. The results of this study could provide researchers with a comprehensive view of research in using of mobile learning in English language learning.

